# The Effects of Personal Pharmacogenetic Testing on the Effects of Pharmacy Student Perceptions of Knowledge and Attitudes Towards Precision Medicine

**DOI:** 10.1101/052043

**Authors:** Dalga Surofchy, Sam S. Oh, Joshua Galanter, Pin Xiang, Megan Li, Su Guo, Tejal Desai, Joseph B. Guglielmo, Kathy Giacomini, Janel Long-Boyle, Alan HB Wu, Esteban G Burchard

**Affiliations:** School of Pharmacy, University of California, San Francisco; Department of Medicine, University of California, San Francisco; Department of Bioengineering & Therapeutic Sciences, University of California, San Francisco; Department of Epidemiology and Biostatistics, University of California, San Francisco; Department of Clinical Pharmacy, University of California, San Francisco; Department of Laboratory Medicine, University of California, San Francisco

**Keywords:** pharmacogenomics, genotyping, pharmacy curriculum, pharmacogenetics, personal pharmacogenetics

## Abstract

**Objective:** To evaluate if pharmacy students’ participation in personal pharmacogenetic (Pgx) testing enhances their knowledge and attitude towards precision medicine (PM).

**Methods:** First-year pharmacy students were offered personalized pharmacogenetic testing as a supplement to a required curricular pharmacogenomics course. Ninety-eight of 122 (80%) students completed pre- and post-course surveys assessing knowledge and attitudes regarding PM; 73 students also volunteered for personal pharmacogenetic testing of the following drug metabolizing enzymes (*CYP2C19*, *CYP2D6*, *UGT1A1*) and pharmacodynamics-relevant proteins (interleukin (IL)-28B & human lymphocyte antigen HLAB*5701).

**Results:** Among the 122 students, we found that incorporating pharmacogenetic testing improved mean knowledge and attitude by 1.0 and 0.3 Likert points, respectively. We observed statistically significant improvements in 100% of knowledge and 70% of attitude-related statements for students who decided to undergo personal pharmacogenetic testing. Students who were enrolled in the course but did not partake in personalized pharmacogenetic testing had similar gains in knowledge and attitude.

**Conclusion:** This study demonstrates the feasibility and importance of educating future pharmacists by incorporating pharmacogenetic testing into professional school curricula. Students who opt not to participate in genotyping may still benefit by learning vicariously through the shared learning environment created by genotyped students.

## INTRODUCTION

Precision Medicine centers around combating human diseases through prevention and treatment based on lifestyle, environment and genetics, a theme embodied by President Obama’s Precision Medicine Initiative.^1^ To increase the acceptance of precision medicine in clinical practice, we must educate future healthcare providers and clinicians. A number of studies^2–10^ regarding the inclusion of pharmacogenomics-related education in professional school curricula suggest promise. Unfortunately, the uptake into U.S. pharmacy programs has been reported as low. In 2010, Murphy et al. surveyed the number of hours U.S. pharmacy schools were dedicating to pharmacogenomics-related education. Only 14.5% of the respondents spent between 31-60 hours on pharmacogenomics-related topics, and 61.3% of respondents described the present state of pharmacogenomics instruction at most schools of pharmacy as “poor.”^11^

Efforts at using genotyping as a teaching technique have been described,^2,4,7–10^ catalyzing the movement towards the adoption of pharmacogenomics into professional school curricula. Our study aims to add to this body of knowledge by providing personal pharmacogenetic testing to first year pharmacy students as an adjunct to a curricular pharmacogenomics course and examining the impact on knowledge and attitude. Innovations in this project include: (1) students having the autonomy to choose the most relevant gene to have genotyped based on their race or personal desire; (2) focusing solely on pharmacogenetic variants, avoiding potential controversy that some direct-to-consumer tests face when providing disease risk assessments (3) utilizing students to lead the initiative.

## METHODS

### Initial Pilot

Prior to our current assessment, a smaller pilot study was conducted among first-year doctor of pharmacy (PharmD) students in Biopharmaceutical Sciences (BPS) 115 (“Genetics and Pharmacogenetics”), a required pharmacogenomics course in the PharmD curriculum. Objectives for BPS 115 were broadly based and derived from components of genetic competencies put forth by the Accreditation Council for Pharmacy Education.^12^ In the spring 2013 offering, twenty-two students volunteered to have their DNA isolated from blood samples and genotyped for variants in *CYP2C19*, a common drug metabolizing enzyme that is important in metabolizing several therapeutic agents, including clopidogrel, a widely used anti-platelet agent. The course directors chose to genotype *CYP2C19* because variants in this gene are known to vary by race, and aside from the ability to metabolize certain drugs, the variants are not known to convey disease risk. This circumvents potential ethical issues that may arise when disclosing disease risk. Some universities offering genomic testing for genetic diseases have been criticized for failing to provide genetic counseling or conducting testing in a non-CLIA-certified laboratory.^13^

The BPS 115 course directors held a lecture session to disclose the results of the students’ genotypes. During this session, course directors reviewed the clinical implications in terms of drug metabolism of different variants of *CYP2C19*. Following the session, students organized a discussion session to ask faculty more questions and create a space for students to continue sharing their learning and genotypic information with other interested classmates. Twenty-two students participated. During this session, students were asked to provide their perceptions/beliefs and attitude regarding the experience, which was recorded in a session transcript taken by a student volunteer and utilized to help develop a formal study protocol. Students unanimously expressed the value of the testing and use of the results as teaching material for the course. Students discussed why it was compelling and crucial to their future as pharmacists and the future of their profession. A sample of representative, unsolicited student comments regarding their experience include:

- “I see personal pharmacogenetic testing in the future of pharmacy. It can be time saving. It is going to be dependent on factors like whether MDs are willing to order genotyping tests instead of starting empirical therapy and dosing, and if we will begin educating our future clinicians. Implementation will require a new generation of MDs/pharmacists to lead this movement.”
- “Information outside of academia regarding pharmacogenomics is limited. Many people in the public are not aware that testing is even available. As leaders/graduates from this university, we have to communicate our knowledge to outside communities and the rest of the world. Having a diverse group of people communicating this information will spread the word about needing research in more ethnically diverse populations. Pharmacists will be the most easily accessible group of healthcare practitioners, so questions about testing will go to us before many in the hospital.”
- “I genuinely enjoyed the class, and I learned a lot. This information inspires me to want to look further into why certain populations are fast metabolizers, or slow metabolizer or do not respond well to certain medications. I would like to personally be involved in pharmacogenomics in the future during my career.”

### Survey Design

Based on feedback from the pilot study, optional personal pharmacogenetic testing was incorporated into BPS 115 the following year. The overall design, implementation and completion of this project was led by three UCSF students; two were members of the School of Pharmacy and one was a graduate PhD student. Their tasks included interpreting feedback from the pilot trial, designing the pre- and post-course surveys, submitting a protocol for approval through UCSF’s institutional review board (IRB) committee, analyzing the data and authoring the manuscript. BPS 115 has been a part of the core UCSF School of Pharmacy curriculum for 12 years. Based on feedback from the pilot trial, the BPS 115 instructors decided to incorporate voluntary personal pharmacogenetic testing during the Spring 2014 term.

A pre-post design was used to evaluate student response. Survey development was informed by the group discussion session following the spring 2013 BPS 115 class. We used students’ responses to identify topics for questions to assess students’ attitudes and knowledge towards precision medicine. A draft of the survey was then piloted among a sample of second-year pharmacy students; these results were used to refine the survey statements. First-year students were excluded from the design process to avoid influencing them during the actual assessment. The final survey included six knowledge statements, ten attitude statements and eight evaluative statements, which appeared only in the post-intervention instrument. (Full instrument available from authors on request.) We chose a Likert-based response format because its common use lends itself to easy understanding by respondents, and answers can easily be quantified and used in statistical tests.

One month before the start of the term, an online Likert survey was administered to 122 first-year UCSF School of Pharmacy (SOP) students enrolled in BPS 115. The survey was re-administered to the same first-year students following completion of the 10-week course. In addition to knowledge- and attitude-assessment statements in the post-course survey, additional evaluative statements were rated, allowing further assessment of students’ opinions about participating in pharmacogenetic testing.

Expected outcomes included: (1) increased understanding of pharmacogenetic concepts in relation to clinical applications, (2) changes in attitudes toward precision medicine and clinical integration of pharmacogenetics, and (3) reports of enhanced learning.

### Survey Administration

The UCSF Committee on Human Research approved the pre- and post-course survey and pharmacogenetic testing. Student coordinators visited the classroom before and after instruction to obtain written consent and email addresses from all students who were interested in participating in the survey. Email addresses were entered into UCSF’s Research Electronic Data Capture (REDCap^14^) system (REDCap, Nashville, Tennessee), a secure online utility for conducting surveys. Once a student logged on to REDCap using their email address, an anonymized, unique identifier was automatically generated and linked to login information. The same identifier was associated with all subsequent surveys, ensuring that no surveys were lost due to individuals forgetting their own self-assigned survey numbers.

While the REDCap survey system collected some basic personal information (e.g., name and UCSF email), it only exported the assigned anonymous identifier with the survey data. Researchers and course faculty members were restricted from accessing the names and email addresses associated with the survey results, helping to ensure that participation was voluntary. In addition, only the researchers were authorized to access the de-identified REDCap data; course directors were not involved in the survey-based assessment.

### Pharmacogenetic Testing

BPS 115 students had the opportunity to volunteer to have their DNA genotyped for several drug-metabolizing enzymes as a “hands-on” personal pharmacogenetic learning experience. Several days were coordinated to collect de-identified saliva samples from students. The samples were analyzed in a UCSF-affiliated CLIA-certified laboratory at San Francisco General Hospital at the rate of $50 per genotype. For the 73 students participating, the total genotyping cost was $3,650, which excludes time donated by laboratory staff to analyze the samples. The UCSF School of Pharmacy covered genotyping costs. Since genotyping was not performed through a commercial supplier, the results were ready sooner than the 3-6 week turnaround time often seen with direct-to-consumer genetic testing companies. The total time student coordinators spent on participant consent and recruitment, DNA collection, and presentation of the data was approximately 140 person-hours. Students were given the option to be genotyped for a gene encoding a drug metabolizing enzyme (*CYP2C19*, *CYP2D6,* or *UGT1A1)* or a pharmacodynamics-relevant protein *(IL28B* or *HLAB*5701).* Each of the genes coding these enzymes/proteins has its own unique clinical implication and varying allele frequency (and therefore varying activity) among ethnic groups (Table 1).

**Table 1:** Drug Metabolizing Enzymes, Function, and Variant Allele Frequencies.

| Enzyme<br>(reference) | Function | Variant Allele<br>Frequency<br>(decreased function)<br>by Race |
| --- | --- | --- |
| CYP2D6<sup>22-24</sup> | Affects large numbers of drugs, notably analgesics, tamoxifen, and antidepressants and medications for attention deficit disorder. | Black: 0-5%<br>Caucasian: 5-14%<br>Asian: 0-1% |
| CYP2C19<sup>23-25</sup> | Affects cardiovascular drugs including clopidogrel and proton pump inhibitors and some antidepressant medications | Black: 5%<br>Caucasian: 2-5%<br>Asian: 19% |
| UGT1A1<sup>26</sup> | Affects some anticancer drugs and is responsible for hyperbilirubinemia induced by Gilbert's syndrome. | Black: 19%<br>Caucasian: 8%<br>Asian: 2% |
| HLA-B*57:01<sup>27</sup> | When present can cause Stevens Johnson Syndrome and delayed hypersensitivity mostly among Asians. | Black: 1%<br>Caucasian: 6-7%<br>Asian: up to 20% |
| IL28B<sup>28</sup> | C/C Alleles predicts drug efficacy towards chronic Hepatitis C infections | Black: 24-50%<br>Caucasian: 8-13%<br>Asian: 0-1% |

Once genotyping was completed, students were given their personal pharmacogenetic information during a regularly scheduled, 120-minute class period for BPS 115. The class session was divided into two parts. During the first part, a pharmacogeneticist was invited to review and discuss each of the genes under evaluation, their variants, clinical significance, and methods for incorporating the information into clinical practice. Five cases were used to illustrate clinical significance of the 5 genotyped genes. Topics included relevant information pertaining to the given gene and relevant drugs, such as irinotecan metabolism (UGT1A1), hepatitis C therapies (IL28b), and Steven Johnson’s syndrome (HLAB*5701). At the end of this discussion, the pharmacogeneticist displayed each of the possible genes via PowerPoint slides and revealed each of the possible variants. Next to each variant was a list of anonymized identifiers so that students were able to privately determine their individual genotype status. The first part of the session was didactic, while the second half of the session emphasized an active learning classroom model. Students were given 15 minutes to discuss the cases presented by the pharmacogeneticist together, and then initiated an open discussion about how different variants may affect pharmacologic or medical management. Students based many of their questions and comments on their personal pharmacogenomic data, as they discussed potential pharmacologic alternatives and interventions (e.g., dose reductions, discontinuation of medications, drug-drug interactions) to account for potential variants. Additionally, students expressed interest in strategies for pharmacists to play a more active role in the future of this specialty. This session did not require specific preparatory work besides attending the course lectures and completing assigned readings^15–17^ pertaining to the course and basic concepts of pharmacogenetics.

### Analysis

We defined our outcome as change in knowledge or attitude regarding precision medicine. Specifically, we assigned integer values to the 5-point Likert scale (i.e., 1 = strongly disagree, 2 = disagree, 3 = neutral, 4 = agree and 5 = strongly agree) and then for each respondent, averaged over all knowledge-related questions, separately for pre- and post-course surveys. This procedure was repeated for attitude-related questions, yielding 4 values for each respondent: pre-course means for knowledge and attitude, as well as post-course means for knowledge and attitude. Thus, our data set was structured in long format as follows: rows represented individual observations—each subject was represented twice (once for pre-course data, and again for post-course data); columns represented responses to individual Likert questions; a separate column represented time (pre-course versus post-course); the last 2 columns represented the mean of knowledge- and attitude-related questions for a given individual (row).

We used linear regression to estimate the effect of participating in BPS 115 on change in mean Likert scores for knowledge and attitude. Specifically, we used our pre/post indicator variable to predict the change in mean knowledge and mean attitude while controlling for sex and race. For example, a pre-survey response of 3 (neutral) followed by a post-survey response of 4 (agree) for mean knowledge would represent a gain of 1 Likert point. This change in mean served as our dependent variable. We assumed that the influence of time between the pre- and post-course surveys was negligible since measurements were made at the same time for all students. Our analysis was stratified into two groups: (1) students who participated in the personal genotyping and the survey (the genotyped group), and (2) those who participated in the survey, but not in personal genotyping (the non-genotyped group). Estimates whose confidence intervals excluded the null value (0) were considered statistically significant at an alpha level of 0.05. Survey results were analyzed using the R statistical programming language (R Core Team, 2015)^18^.

## RESULTS

In total, 98 (80%) of the 122 students enrolled in the spring 2014 BPS 115 course voluntarily completed the pre- and post-course surveys. Of these 98, 73 (74.5%) students also took part in genotyping, leaving 25 students (25.5%) to comprise the surveyed but not genotyped group. Selected demographic characteristics of the students are summarized in Table 2. Baseline scores in knowledge and attitude were similar for both groups. The mean baseline Likert score for knowledge statements was (3.03) in the genotyped group and (3.14) in the non-genotyped group. For the attitude statements, the mean baseline score was (3.85) in the genotyped group and (3.83) in the non-genotyped group.

**Table 2:**
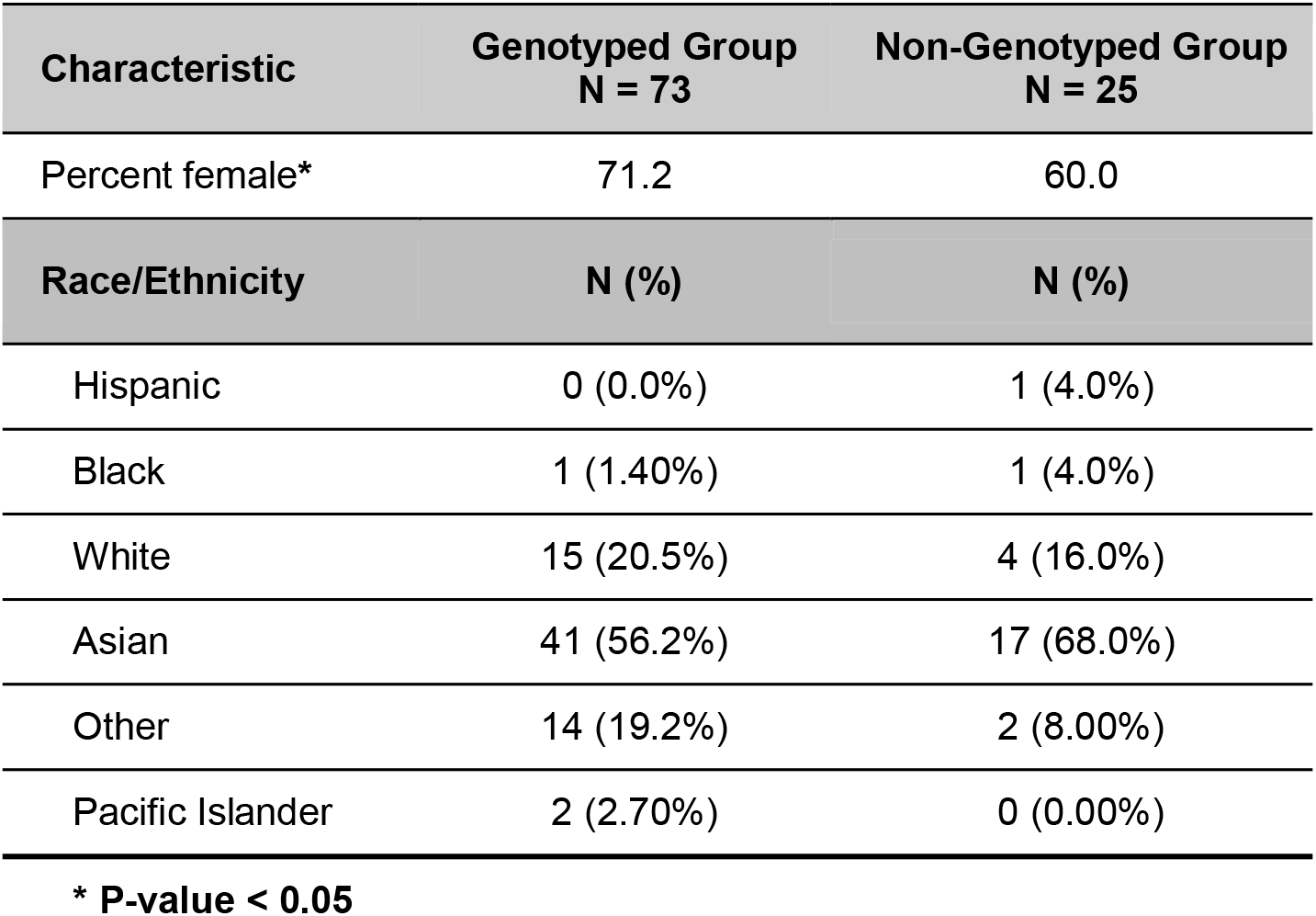
Gender and Race/Ethnicity of Participants and Nonparticipants.

The results for change in mean knowledge and mean attitude for all students combined, and stratified by genotyping status, are summarized in Figure 1. By the end of the course, statistically significant increases were observed for mean knowledge (+1.0 Likert points) and mean attitude (+0.3 Likert points). The gain in mean knowledge was not significantly different between the genotyped (0.99) and non-genotyped (1.04) groups (p = 0.77). While the gain in mean attitude was slightly higher among those who did not participate in genotyping (0.34) versus those who did (0.29), the difference was not statistically significant (p = 0.74). The correlation between pre- and post-survey responses was fair for mean knowledge (Pearson’s r = 0.50) and mean attitude (r = 0.46).

**Figure 1:**
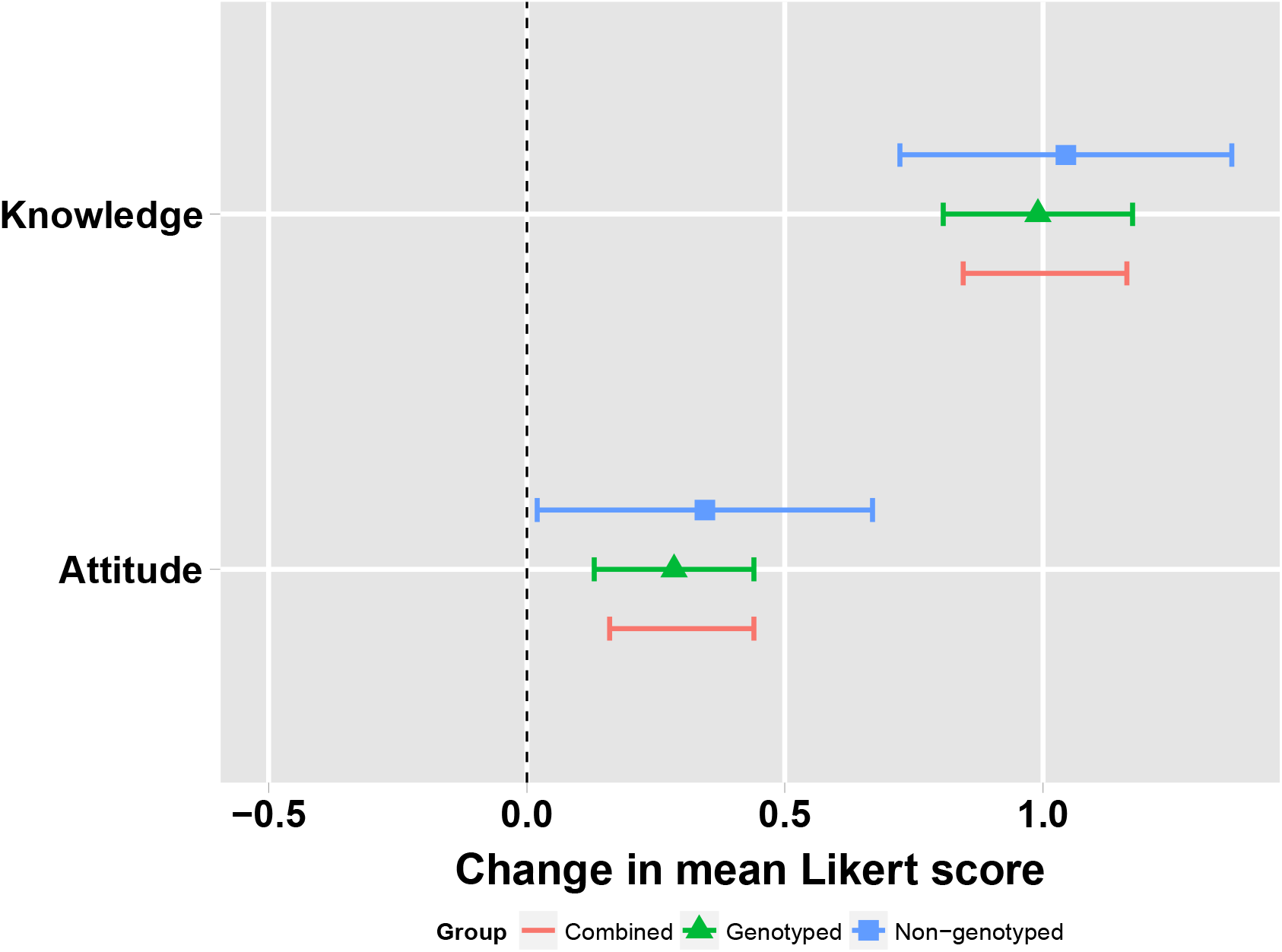
Overall Mean Change in Knowledge and Attitude.

We also asked students to evaluate their experiences. Table 3 reports responses from the genotyped and non-genotyped groups. Changes in knowledge and attitude for specific questions are summarized in Table 4.

**Table 3.** Responses to Evaluative Statements from Genotyped and Non-Genotyped Groups.

| **Genotyped Group N = 73** |  |  |  |  |  |
| --- | --- | --- | --- | --- | --- |
| **Question** | **Strongly Disagree**<br>**N (%)** | **Disagree**<br>**N (%)** | **Neutral**<br>**N (%)** | **Agree**<br>**N (%)** | **Strongly Agree**<br>**N (%)** |
| Glad to have participated in Pgx testing | 0 (0%) | 2 (3.00%) | 6 (8.00%) | 31 (42.0%) | 34 (47.0%) |
| I believe I have a better understanding of Pgx principles | 0 (0%) | 1 (1.00%) | 10 (14.0%) | 37 (51.0%) | 25 (34.0%) |
| Felt more engaged because I had undergone Pgx testing | 1 (1.00%) | 2 (3.00%) | 14 (19.0%) | 26 (36.0%) | 30 (41.0%) |
| My participation reinforced concepts taught in BPS 115 | 1 (1.00%) | 1 (1.00%) | 10 (14.0%) | 34 (47.0%) | 27 (37.0%) |
| **Non-Genotyped Group N = 25** |  |  |  |  |  |
| **Question** | **Strongly Disagree**<br>**N (%)** | **Disagree**<br>**N (%)** | **Neutral**<br>**N (%)** | **Agree**<br>**N (%)** | **Strongly Agree**<br>**N (%)** |
| I regret not participating in Pgx | 0 (0%) | 5 (20.%) | 5 (20.0%) | 11 (44.0%) | 4 (16.0%) |
| Classmates' Pgx results reinforced concepts | 0 (0%) | 1 (4.00%) | 7 (28%) | 14 (56.0%) | 3 (12.0%) |
| I am interested in participating in more comprehensive genetic tests | 0 (0%) | 1 (4.0%) | 7 (28%) | 13 (52.0%) | 4 (16%) |
| If offered again, I would undergo Pgx testing | 0 (0%) | 2 (8.00%) | 8 (32%) | 9 (36.0%) | 6 (24.0%) |

BPS 115: Biopharmaceutical sciences 115, "Genetics and Pharmacogenetics"
Pgx: Pharmacogenetics

**Table 4:** Knowledge and Attitude Assessment Questions from Pre and Post Survey.

|  | **Genotyped (N=73)**<br>**Effect Size; 95% CI** |  | **Non-Genotyped (N=25)**<br>**Effect Size; 95% CI** |  |
| --- | --- | --- | --- | --- |
|  | **Effect Size** | **95% CI** | **Effect Size** | **95% CI** |
| **Knowledge** |  |  |  |  |
| **Change in mean knowledge** | **0.99** | **0.81-1.17** | **1.04** | **0.72-1.37** |
| 1. I understand what precision medicine is. | 0.64 | 0.42-0.87 | 0.84 | 0.41-1.27 |
| 2. I understand what a pharmacogenetic test is. | 0.89 | 0.67-1.11 | 1.20 | 0.78-1.62 |
| 3. I understand how a pharmacogenetic test differs from a genetic test for disease risk. | 1.18 | 0.92-1.44 | 0.96 | 0.42-1.50 |
| 4. I am aware of the types of knowledge and resources needed to interpret a pharmacogenetic test result. | 1.32 | 1.05-1.58 | 1.08 | 0.59-1.57 |
| 5. I understand how to evaluate the clinical validity and utility of a pharmacogenetic test. | 1.19 | 0.91-1.47 | 1.32 | 0.86-1.78 |
| 6. I understand the risks, benefits, and ethical considerations of personal genetic testing. | 0.71 | 0.47-0.95 | 0.84 | 0.42-1.26 |
| **Attitude** |  |  |  |  |
| **Change in mean attitude** | **0.29** | **0.13-0.44** | **0.34** | **0.02-0.67** |
| 1. The use of personal genetic information in health care is beneficial to patients. | 0.29 | 0.08-1.25 | 0.20 | -0.22-0.62 |
| 2. The use of personal genetic information in health care may cause unnecessary harm to patients. | -0.01 | -0.32-0.31 | 0.18 | -0.42-0.77 |
| 3. In addition to factors like age, race, and drug interactions, genetic information is an important consideration during routine clinical practice. | 0.28 | 0.05-0.48 | 0.28 | -0.13-0.69 |
| 4. I would recommend pharmacogenetic testing for a patient. | 0.49 | 0.26-0.65 | 0.48 | 0.01-0.95 |
| 5. I would recommend pharmacogenetic testing for a family member. | 0.47 | 0.25-0.62 | 0.44 | -0.03-0.91 |
| 6. Pharmacogenetic testing, when applicable, should be integrated into patient care. | 0.47 | 0.26-0.70 | 0.50 | 0.09-0.90 |
| 7. Pharmacists should be trained to | 0.26 | 0.06-0.48 | 0.44 | 0.08-0.80 |
| interpret and apply pharmacogenetic test results. |  |  |  |  |
| 8. Pharmacogenetics should be integrated into the curricula at all pharmacy schools. | 0.34 | 0.12-0.55 | 0.48 | -0.04-0.92 |
| 9. Pharmacogenetics will likely play an important role in my future career. | 0.09 | -0.15-0.29 | 0.28 | -0.17-0.73 |
| 10. Pharmacists play a crucial role in the future of precision medicine. | 0.18 | -0.02-0.38 | 0.24 | -0.17-0.65 |

## DISCUSSION

We found that incorporating genetic testing as an adjunct to the School of Pharmacy PharmD curriculum enhanced students’ self-reported knowledge and attitudes of precision medicine. We observed statistically significant increases across all knowledge assessment statements after the study. This finding provides support that an interactive hands-on approach to educating future pharmacists about pharmacogenetics is a curricular change that could benefit professional doctorate programs.

Previous pharmacogenetics education efforts in schools of pharmacy^4,7,9,10^ and medicine^2^ have influenced our approach. Most recently in 2016, Adams et al.^10^ published results from the University of Pittsburgh demonstrating significant improvements in PharmD students’ knowledge and attitude after participating in personal pharmacogenetic testing through a commercial genotyping service. Since utilizing commercial services raised concerns about incidental findings of disease risk, we decided to eliminate this risk by focusing on genes that were solely pharmacogenetically relevant. Additional insight from the aforementioned literature that guided our approach included ensuring an interactive and hands-on approach^2^. For example, we sought to increase relevance and student interaction by allowing the selection of pharmacogenetic genes that may have been important to our students based on their ethnicity or personal interests.

Our curricular approach to increasing knowledge and improving attitudes towards pharmacogenetic testing and precision medicine yielded encouraging results. The majority (80%) of students completed pre- and post-course surveys, and 74.5% took part in personal pharmacogenetic testing. Based on our experience, implementing personal pharmacogenetic testing in U.S. pharmacy school curricula need not be extremely arduous. Student participation was very high in the absence of incentives; the time and effort dedicated toward collection and processing of DNA was fairly minimal; performing genotyping in-house was efficient and allowed for exclusion of disease-associated variants; and the discussion of genotyping results was limited to only one class session. Instructors could limit their selection of genetic tests to inexpensive ones to optimize widespread dissemination of an educational session of this type.

One of our most noteworthy findings was in regard to Knowledge Statement #4 (Table 4): “I am aware of the types of knowledge and resources needed to interpret a pharmacogenetic test.” The reported change was fairly large among genotyped students, (1.32, 95%CI: 1.05-1.58), demonstrating that students felt confident utilizing their resources to interpret a pharmacogenetic test. This observation suggests that a curriculum designed to include similar personal pharmacogenetic testing may help prepare students to keep up with the precision medicine revolution.

The gain in knowledge and attitude for both the genotyped and non-genotyped group is interesting and questions whether our results are attributable to personal pharmacogenetic testing versus traditional didactic coursework. The difference in improvement between groups is greater for the knowledge-related items than for attitude, consistent with the belief that knowledge affects attitude, which in turn affects behavior.^19,20^ The genotyped group had tighter confidence intervals and more robust data as we would have expected based on sample size. However, that both groups improved in knowledge and attitude is encouraging, suggesting that once participation in genetic testing surpasses some threshold, non-genotyped students may learn vicariously through experiences and learning environment created by genotyped students. In fact, 68% of non-genotyped students agreed or strongly agreed that their classmates’ participation in genotyping positively impacted their learning in the course (Table 3). In essence, the rising tide of pharmacogenetic education lifted all boats.

Potential biases and limitations should be considered when reviewing the results of our study. Knowledge and attitude were measured by self-assessment. This method is not as robust as objective data (e.g., exam questions). Some unmeasured characteristics of genotyped students (e.g., attitudes toward providing biological samples) may have differed from non-genotyped students such that comparison of pre-post results between these groups could have been biased. However, we found no significant difference in baseline knowledge or attitude between the two groups. Given that the reported change for all knowledge and 9/10 of the attitude assessment statements were positive, regardless of genotyping status, we feel that the influence of this type of selection bias was minimal. Results for the non-genotyped group may have been underpowered given the smaller number of students who chose not to be genotyped (25 versus 73). Our analyses were conducted under the assumption that the intervals between Likert values are equal. We felt it reasonable, for example, to assume that “Agree” is halfway between “Neutral” and “Strongly agree,” a common assumption practiced in analysis of survey results.^21^ Isolating the impact of genetic testing through an experimental design without overly disrupting the course structure posed logistical challenges. As with intervention trials, preventing crossover (e.g., non-compliance or contamination) between treatment groups would have been difficult—a common case in education-related studies.

To overcome these limitations, a future cluster-based randomized study with several pharmacy school curricula could be implemented. Since self-efficacy—which was not measured in this study—is useful for predicting future behavior, we plan to contact these students for a long-term follow-up to assess the lasting effects that personal pharmacogenetic testing has had on their personal and professional lives.

## SUMMARY

First-year PharmD students who volunteered to participate in personal pharmacogenetic testing showed statistically significant increases in self-reported knowledge and attitudes towards precision medicine. This increase was also observed among students who were enrolled in the course, but did not partake in personalized pharmacogenetic testing, likely a result of engagement with their classmates and faculty. This dynamic allows room for pharmacy schools to personalize the incorporation of pharmacogenetic testing into their curricula.

Our study was comprised of three innovative elements: (1) providing students the autonomy to choose the most relevant gene for genotyping; (2) focusing solely on pharmacogenetic variants, avoiding potential controversy associated with some direct-to-consumer tests that also assess disease risk; and (3) utilizing students to lead the initiative. These three components may aid dissemination of similar projects at other institutions.

## ACKNOWLEDGEMENTS

We are grateful to Sandra Salazar and the UCSF School of Pharmacy for the support and contributions toward the IRB process.

